# Visual effort moderates postural cascade dynamics

**DOI:** 10.1101/2020.07.17.209486

**Authors:** Madhur Mangalam, I-Chieh Lee, Karl M. Newell, Damian G. Kelty-Stephen

## Abstract

Standing still and focusing on a visible target in front of us is a preamble to many coordinated behaviors (e.g., reaching an object). Hiding behind its apparent simplicity is a deep layering of texture at many scales. The task of standing still laces together activities at multiple scales: from ensuring that a few photoreceptors on the retina cover the target in the visual field on an extremely fine scale to synergies spanning the limbs and joints at smaller scales to the mechanical layout of the ground underfoot and optic flow in the visual field on the coarser scales. Here, we used multiscale probability density function (PDF) analysis to show that postural fluctuations exhibit similar statistical signatures of cascade dynamics as found in fluid flow. In participants asked to stand quietly, the oculomotor strain of visually fixating at different distances moderated postural cascade dynamics. Visually fixating at a comfortable viewing distance elicited posture with a similar cascade dynamics as posture with eyes closed. Greater viewing distances known to stabilize posture showed more diminished cascade dynamics. In contrast, nearest and farthest viewing distances requiring greater oculomotor strain to focus on targets elicited a dramatic strengthening of postural cascade dynamics, reflecting active postural adjustments. Critically, these findings suggest that vision stabilizes posture by reconfiguring the prestressed poise that prepares the body to interact with different spatial layouts.

## 1. Introduction

Standing still and visually fixating on a target in front of us is a seemingly simple task. However, hiding within this apparent simplicity is a deep layering of texture at many scales. On an extremely fine-scale, task completion depends only on ensuring that a few photoreceptors on the retina cover the entire target in the visual field. Standing still and visually fixating at much coarser scales rests on the mechanical layout of the ground underfoot and optic flow in the visual field before you. Between optic flow and mechanical layout, standing still relies on synergies spanning the limbs and joints at smaller scales and subtle postural perturbations, triggering reflex arcs and muscle twitches, at yet smaller scales. The task of standing still and visually fixating laces together activities at these multiple scales, entailing a close connection among events of widely varying scales, from whole-body postural fluctuations at large scales, to head sway and eye movements at medium scales, and photoreceptor activity at the finest scales [1–3]. Consequently, the postural center of pressure (CoP) fluctuations—which directly relate to sway—exhibit similar statistical signatures of cascade dynamics as found in fluid flow [4]. The metaphor of ‘cascades,’ as in tumbling water that accelerates and splashes down a rockface, captures some critical aspects of the dynamics of emergent behavior [5–7]. Destabilizing posture emphasizes the postural cascade dynamics [8,9]. Here, we propose that perturbing the visual system has a similar effect of strengthening postural cascade dynamics, in which the optical flow varies with fixation distance, as a meandering stream joins other streams and gathers momentum into rushing rapids, becoming more turbulent along the way. Whereas multiscale entropy addresses the difference of variation across different scales [10,11], cascade dynamics explicitly implicates nonlinear interactions lacing these scales together.

Visually guided posture presents a curious case in which light entails mechanical change in bodily center of mass [12–17]. Beyond minuscule photoreceptor responses, large-scale layout of visible surfaces supports optic flow with bodily sway. Optic flow is the expansion, contraction, or rotation of visible surfaces as the body moves forward or backward, or turns to the side, respectively. Relative accelerations of visible surfaces provide the standing participant with rich information about the layout of objects. Placing a fixation target at different viewing distances perturbs the optic flow: changing the relative accelerations of visible surfaces as well as imposing retinal-specific constraints of oculomotor convergence (i.e., higher oculomotor strain in specific or visual effort in general) [18,19], prompting new head- and torso-sway (Fig. 1). Indeed, optic flow relies on nesting of behaviors across the aforementioned range of scales, including the organism nested in the task environment at the largest scale right down to the hierarchical organization of photoreceptor response of optic flow at the finest scale [3]. Comfortable viewing distances (50-100cm) requiring least oculomotor strain should most resemble posture with eyes closed [20,21]. Medium viewing distances beyond most-comfortable viewing in the range of 200 cm stabilize posture [22]. Extremely distant or extremely near targets will be harder to focus on and have a much slower or faster optic flow, respectively, requiring more sway to resolve the image [23].

**Fig. 1.**
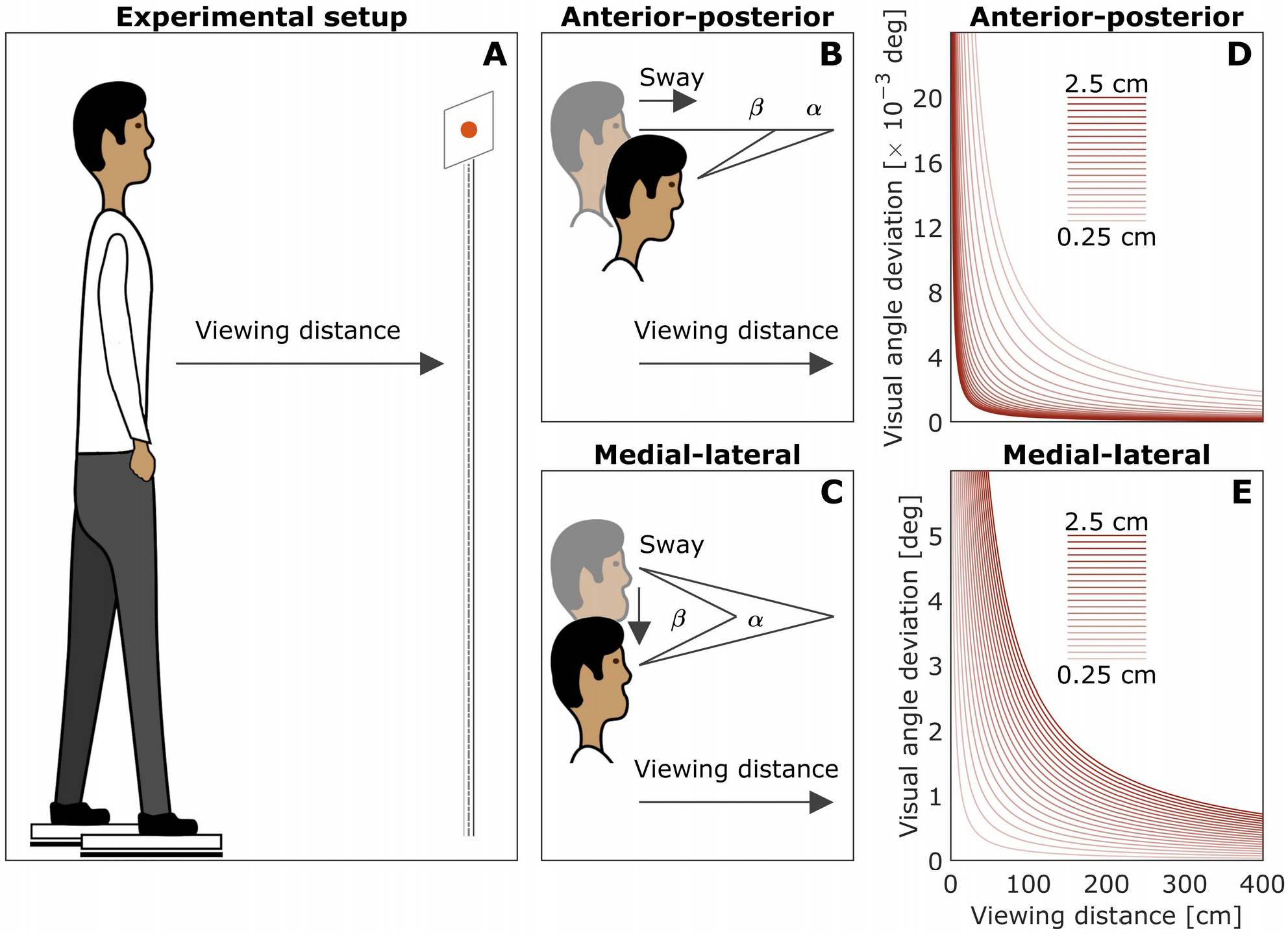
Schematic of the task and effects of eye-to-target distance on postural sway. (*A*) The suprapostural viewing task of standing quietly with the eyes fixated at a distant visual element. (*B* and *C*) Visual angle gain for short vs. long eye-to-target distances along the anterior-posterior (AP) and medial-lateral (ML) axes. (*D* and *E*) Visual angle gain as a function of eye-to-target distance for different sway magnitudes. Closer targets increase *AP* sway, whereas farther targets increase *ML* sway.

**Fig. 2.**
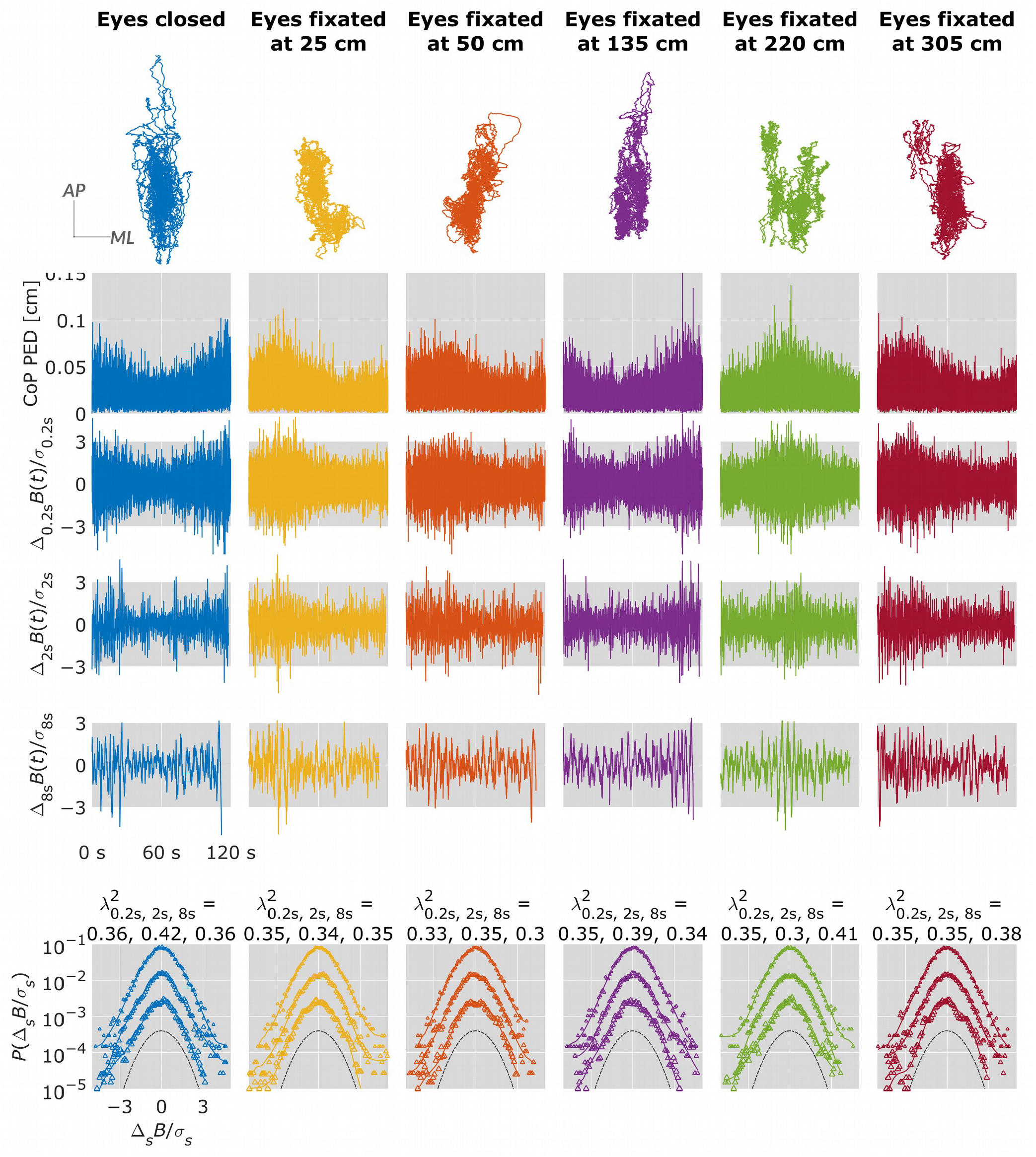
Multiscale PDF characterization of postural fluctuations in a representative participant maintaining quiet stance for 120 s with eyes closed, and eyes fixated at a point at a distance of 25, 50, 135, 220, and 305 cm from the eyes. From to bottom: CoP trajectories along the anterior-posterior (AP) and medial-lateral (ML) axes. CoP PED series. {Δ_*s*_ *B*(*i*)} for *s* = 0.5, 2, and 8 s. Standardized PDFs (in logarithmic scale) of {Δ_*s*_ *B*(*i*)} for *s* = 0.5, 2, and 8 s (from top to bottom), where *σ*_*s*_ denotes the *SD* of {Δ_*s*_ *B*(*i*)} for *s* = 0.5, 2, and 8 s (from top to bottom), where *σ*_*s*_ denotes the *SD* of {Δ_*s*_ *B*(*i*)}.

Testing these hypotheses requires examining the divergence of postural fluctuations from Gaussian form. Gaussian distributions are classically the result of adding very many independent (i.e., non-interacting) variables. Interactivity amongst the participating variables promotes persistent if also intermittent, uneven growth of variance with timescale. Under the central limit theorem, homogeneity of variance keeps measurements within short, thin tails of a Gaussian distribution. But intermittent growth in variance manifests in distributions with longer, heavier tails specifically due to nonlinear temporal correlations [24,25]. Testing these tails is even more difficult because they are by definition only populated by extreme and relatively rare values. However, fortunately, hydrodynamic-cascade research has yielded an analytical method for estimating non-Gaussianity based on querying better-populated regions of the distribution [26]. Applying this method to physiological measures produces estimates of non-Gaussianity that predict clinical outcomes [27–31] and that change with variation in both endogenous postural control (i.e., unmanipulated factors that respond to but also exert effects upon other variables of the postural system) and exogenous postural demands [8,9]. Here, we used this method to investigate how constraints imposed by visual effort affect postural cascade dynamics. Participants were asked to minimize postural sway while maintaining quiet stance for 120 s in six different viewing conditions: eyes closed and eyes fixated on a point on a screen at a distance of 25, 50, 135, 220, and 305 cm from the eyes. We predicted that postural cascade dynamics while fixating at a 50-cm distance would resemble eyes-closed posture (Hypothesis-1), and that postural cascade dynamics would be weakest while fixating at medium (i.e., 130- and 220-cm) distances (Hypothesis-2) but strongest at extremely near (i.e., 25-cm) and extremely far (i.e., 305-cm) distances (Hypothesis-3).

## 2. Materials and methods

The present study is a reanalysis of data from a previous study examining postural control in different viewing conditions [32].

### 2.1. Participants

Seven men and eight women (18–40 years old) participated after providing institutionally-approved informed consent.

### 2.2. Experimental setup and procedure

Each participant stood barefoot on two force plates (AMTI Inc., Watertown, MA) recording 3D moments and ground reaction forces at 100 Hz—one foot on each plate, 25 cm apart, before a white background spanning his/her visual field (Fig. 1A). A square white screen (5×5 cm) mounted on a tripod was placed at specific distances in front of the participant. From behind the participant, a laser pen projected a static light point on the screen’s center.

Each participant experienced six different viewing conditions (eyes-closed and eyes fixated on the screen’s light point at a distance of 25, 50, 135, 220, and 305 cm in the front), each three times in randomized order over a single 90-min session. In each trial, each participant was instructed to minimize the postural sway by maintaining visual fixation at the light point projected on the screen for 120 s. Each participant had a 5-min break for rest after every six trials.

### 2.3. Data processing

All data processing was performed in MATLAB 2019b (Matlab Inc., Natick, MA). Trial-by-trial ground reaction forces yielded a 2D center of pressure (CoP) series, each dimension describing anterior-posterior (AP) and medial-lateral (ML) axes. Each 60-s trial yielded 12000-sample 2D CoP series and 11999-sample 2D CoP displacement series. Finally, a 1D CoP sample-to-sample planar Euclidean displacement (PED) series described postural fluctuations along the transverse plane. Iterated Amplitude Adjusted Fourier Transformation (IAAFT) provided phase-randomized surrogates using original series’ spectral amplitudes to preserve only linear temporal correlations [33].

### 2.4. Canonical indices of endogenous postural fluctuations

Three linear indices were computed for each CoP PED series: (i) Mean of all fluctuations (CoP_PED_*Mean*), (ii) Standard deviation (CoP_PED_*SD*), and (iii) Root mean square (CoP_PED_*RMSE*).

Four nonlinear indices were computed for each CoP PED series: (i) Sample entropy (CoP_PED_*SampEn*), indexing the extent of complexity in the signal [34], using *m* = 2 and *r* = 0.2 [35]. (ii) Multiscale sample entropy (CoP_PED_*MSE*), indexing signal complexity over multiple timescales [10], using *m* = 2, *r* = 0.2, and *τ* = 20 [35]. (iii) We used detrended fluctuation analysis [36] to compute Hurst’s exponent, *H*_fGn_, indexing temporal correlations in original CoP PED series (CoP_PED_*H*_fGn_Original_) and shuffled versions of each series (CoP_PED_*H*_fGn_Shuffled_) using scaling region: 4, 8, 12,… 1024 [37]. Shuffling destroys the temporal structure of a signal. (iv) Multifractal spectrum width, Δ*α*, indexing the extent of multifractal temporal correlations in the signal. Chhabra and Jensen’s direct method [38] was used to compute Δ*α* for original CoP series (CoP_PED_Δ*α*_Original_) and IAAFT surrogate (CoP_PED_Δ*α*_Surrogate_) [6].

### 2.5. Multiscale probability density function (PDF) analysis

Multiscale PDF analysis characterizes the distribution of abrupt changes in CoP PED series {*b*(*t*)} using the PDF tail. The first step is to generate {*B*(*t*)} by integrating {*b*(*t*)} after centering by the mean *b*_*ave*_ (Fig. S1A):

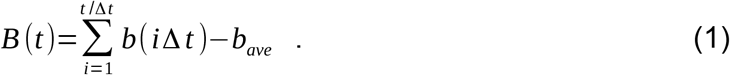

A a 3^rd^ order polynomial detrends {*B* (*t*)} within *k* overlapping windows of length 2*s, s* being the timescale (Fig. S1B). Intermittent deviation Δ_s_*B*(*t*) in *k*^*th*^ window from 1+*s* (*k* −1) to *sk* in the detrended time series {*B*^*d*^(*t*)=*B*(*t*)−*f*_*fit*_(*t*)} is computed as Δ_s_*B*^*d*^ (*t*)=*B*^*d*^ (*t* + *s*)−*B*^*d*^(*t*), where 1+*s* (*k* −1)≤*t* ≤*sk* and *f*_fit_ (*t*) is the polynomial representing the local trend of {*B* (*t*)}, of which the elimination assures the zero-mean probability density function in the next step (Fig. S1C). Finally, Δ_*s*_*B* is normalized by the SD (i.e., variance is set to one) to quantify the PDF.

To quantify the non-Gaussianity of Δ_*s*_*B* at timescale *s*, the standardized PDF constructed from all the Δ_*s*_*B*(*t*) values is approximated by the Castaing model [26], with *λ*_*s*_ as a single parameter characterizing the non-Gaussianity of the PDF. *λ*_*s*_ is estimated as

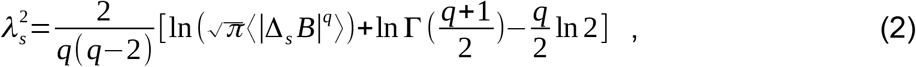

where ⟨|Δ_s_*B*|^*q*^ ⟩ denotes an estimated value of *q*^*th*^ order absolute moment of {Δ_s_*B*}. As 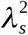 increases, the PDF becomes increasingly peaked and fat-tailed (Fig. S2A). 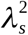 can be estimated by Eq. (2) based on *q*^*th*^ order absolute moment of a time series independent of *q*. Estimating 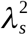 based on 0.2^th^ moment (*q* = 0.2) emphasizes the center part of the PDF, reducing the effects of extreme deviations due to heavy-tails and kurtosis. We used 0.2^th^ moment because estimates of 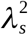 for a time series of ∼ 12000 samples are more accurate at lower values of *q* [39].

Cascade-type multiplicative processes yield the inverse relationship 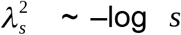 (Fig. S2B) [29]. For the present purposes, we quantified 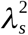 for each original CoP PED series and corresponding IAAFT surrogate at timescales 5 to 1000 samples (i.e., 50 ms to 10 s) at steps of 5 samples (50 ms).

We used Akaike Information Criterion (AIC) weights obtained via the maximum likelihood estimation (MLE) to determine whether the PDF at each timescale was a power law, lognormal, exponential, or gamma distribution. We include this analysis also to show how AIC-based MLE sensitivity to lognormal-like heavy tails differs from multiscale PDF sensitivity to the bulk of the distribution.

### 2.6. Statistical analysis

A linear mixed-effect (LME) model using lmer [40] in R package *lme4 tested* 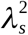 vs. log-timescale curves for orthogonal linear and quadratic polynomials, for interactions with grouping variables (Viewing condition × Original, where Original encoded differences in original series from surrogates) and with indices of endogenous postural fluctuations (Section 2.4.). Statistical significance was assumed at the alpha level of 0.05 using R package *lmerTest* [41]. To test how lognormality changed with log-timescale, a generalized linear mixed-effect (GLME) fit changes in Lognormality as a dichotomous variable using orthogonal linear, quadratic, and cubic polynomials and tested interaction effects of grouping variables (Viewing condition × Original) with those polynomials using glmer [42] in *lme4*.

## 3. Results

The eyes-closed condition elicited nonlinearity in non-Gaussianity with log-timescale, showing a stronger linear increase in 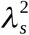 up to the timescale of roughly 75 samples or 750 ms (*B* = 0.95, *P* = 0.031) and a stronger negative quadratic, downward-facing parabolic form (*B* = –3.78, *P* < 0.001) for the original CoP PED series than the surrogates (Table S1; Fig. 3A). Fixating at 25 and 305 cm elicited a stronger linear increase (*B* = 2.43, *P* < 0.001; and *B* = 3.09, *P* < 0.001, respectively) in the 25- and 305-cm viewing conditions while reversing the downward-facing parabolic form of the eyes-closed condition more weakly (*B* = 4.42, *P* < 0.001; and *B* = 1.85, *P* = 0.003, respectively) of 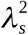 vs. log-timescale curves (Figs. 3B and 3F). Fixating the eyes at 135 and 220 cm elicited a reversal of linear (*B* = –2.98, *P* < 0.001; and *B* = –2.32, *P* < 0.001, respectively) and downward-facing parabolic forms (*B* = 1.81, *P* = 0.004; and *B* = 2.00, *P* = 0.001, respectively) of 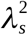 vs. log-timescale curves in Figs. 3D and 3E. In contrast, fixating at 50 cm neither affected the linear form (*B* = –0.09, *P* = 0.879) nor the downward-facing parabolic form (*B* = 0.73, *P* = 0.244; Fig. 3C). Thus, the 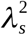 vs. log-timescale curves differ between the eye-closed condition and all the eyes-open conditions except the eyes-fixated-at-50-cm condition. Furthermore, whereas the 50- and 305-cm viewing conditions *show significant growth of the* 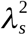 vs. log-timescale curves, the 135- and 220-cm viewing conditions exhibit significant decay of this curve with scale.

**Fig. 3.**
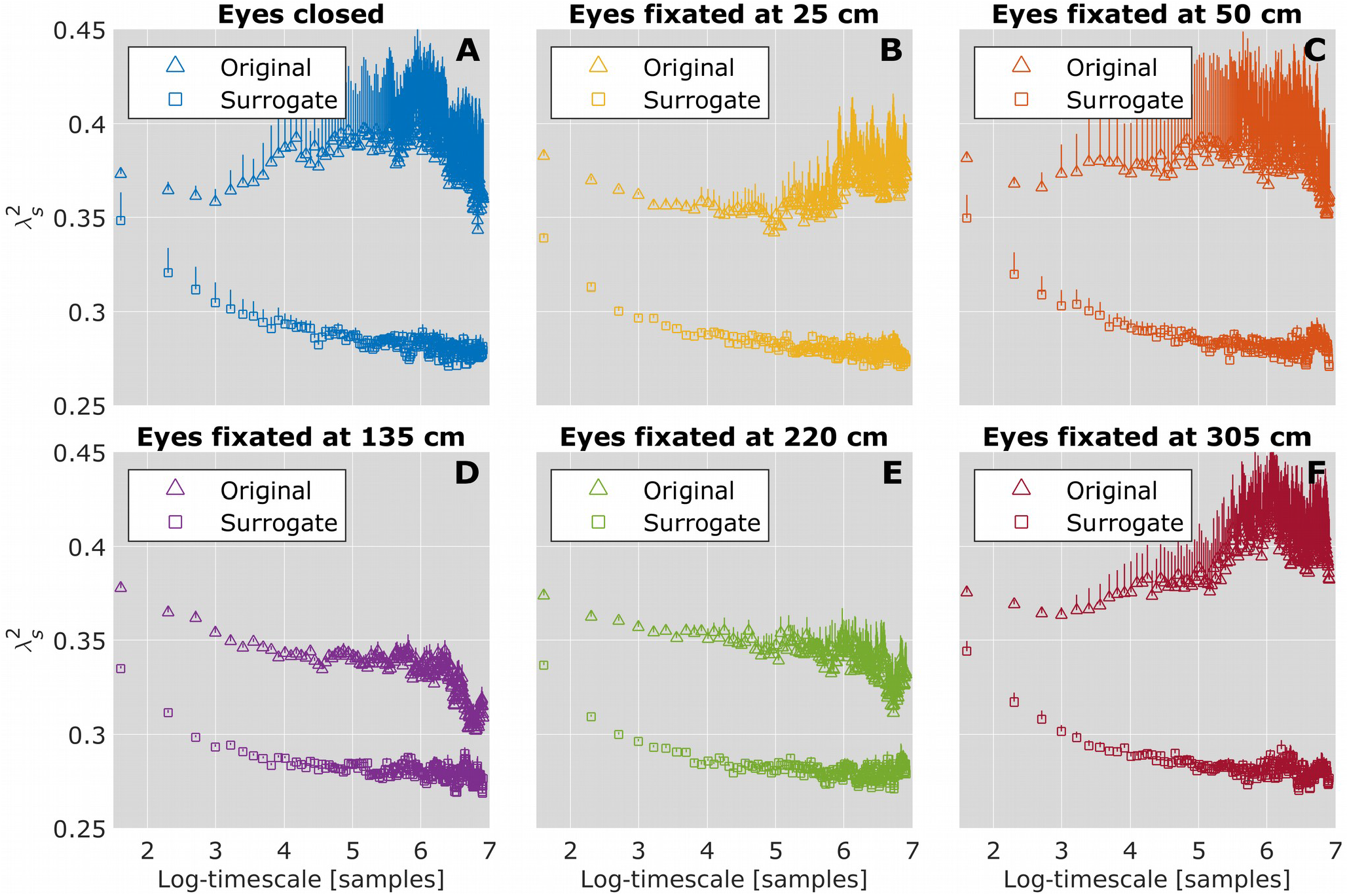
Log-timescale dependence of the non-Gaussianity index 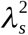. Mean values of 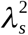 for the participants standing quietly for 120 s in different viewing conditions. (A) Eyes closed. (B) Eyes fixated at 25 cm. (C) Eyes fixated at 50 cm. (D) Eyes fixated at 135 cm. (E) Eyes fixated at 220 cm. (F) Eyes fixated at 305 cm. Vertical bars indicate ±1*SEM* (*N* = 15). Notice that *the* 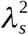 vs. log-timescale curves differ between the eye-closed condition all the eyes-open conditions except the eyes-fixated-at-50-cm condition.

All indices, except sample entropy of postural fluctuations, showed predictive effects on 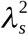 · 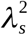 decreased with increases in root mean square, and increased with increases in mean, standard deviation, and multiscale entropy of postural fluctuations (Table S2). Nonetheless, greater ‘original-minus-shuffled/surrogate’ differences in fractality and multifractality diminished 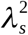. Hence, evidence of multifractality with weak difference from the surrogates suggests heavy-tailed distributions and thus non-Gaussianity [30,31].

Interactions of these indices with linear and quadratic terms in the first model significantly improved the model fit (*χ*_315_ = 153961, *P* < 0.001; Table S2). Most variability in the quadratic form of the 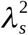 vs. log-timescale curves depended on these indices, as all but two of 41 significant (*P* < 0.05) quadratic effects relate to interactions of these indices with log-timescale. That is, the parabolic tempering of non-Gaussianity across viewing conditions depended on endogenous postural control. A cubic polynomial improved model fit (*χ*^2^(120) = 2061.8, *P* < 0.001), reflecting 17 significant interactions with the cubic effect, but left all lower-order terms unchanged in coefficients and significance (Table S3).

Postural instability typically weakens the growth of lognormality in postural fluctuations with log-timescale [8,9]. The original CoP PED series exhibited less lognormality than the surrogates (*B* = –4.28, *P* < 0.001; Table S4; Fig. 4). Lognormality grew with log-timescale: a linear increase (*B* = 2219.40, *P* = 0.016), quadratic decrease (*B* = –1678.38, *P* = 0.052), and cubic increase (*B* = 559.57, *P* = 0.051) in the original CoP PED series than the surrogates. However, polynomial profiles of the growth of lognormality with log-timescale did not differ among the conditions. Hence, observed differences in the 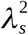 vs. log-timescale curves reflect differences in the bulk of the postural-fluctuations distributions rather than in the tail.

**Fig. 4.**
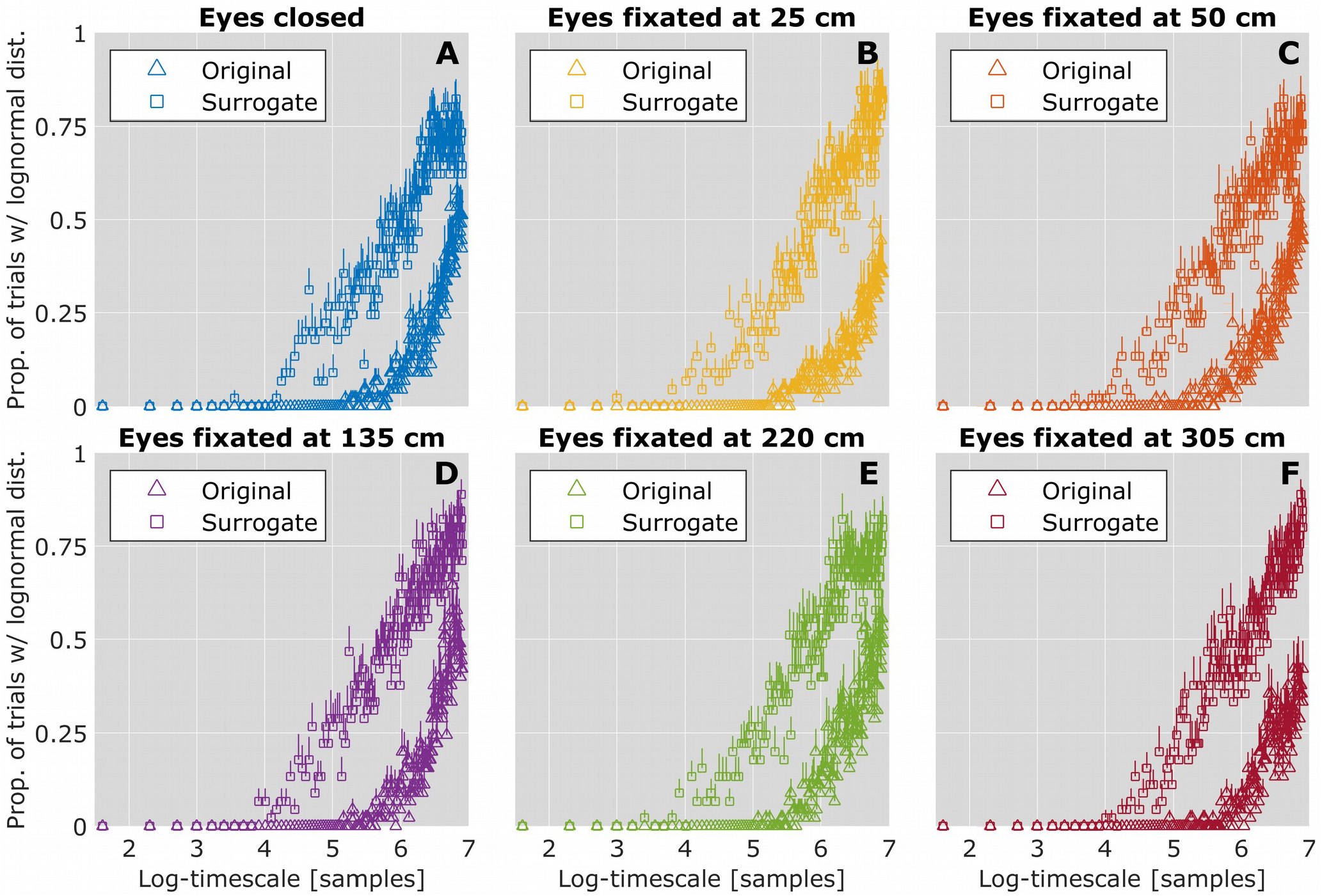
Log-timescale dependence of the mean proportion of trials with a lognormal distribution for the participants standing quietly for 120 s in different viewing conditions. (A) Eyes closed. (B) Eyes fixated at 25 cm. (C) Eyes fixated at 50 cm. (D) Eyes fixated at 135 cm. (E) Eyes fixated at 220 cm. (F) Eyes fixated at 305 cm. Vertical bars indicate ±1*SEM* (*N* = 15). Notice that the polynomial profiles of the growth of lognormality with log-timescale do not differ between the eyes-closed condition and any of the eyes-open conditions.

## 4. Discussion

Visual effort moderates postural cascade dynamics, given that standing still and visually fixating at shorter and longer distances unleashed stronger cascade dynamics than standing still (Hypothesis-3) and visually fixating at an optimal distance or standing still without visually fixating (i.e., with eyes closed; Hypotheses-1 & 2). Importantly, these effects are not reducible to the mere availability of visual information, as standing still and visually fixating at an optimal distance and standing still with eyes closed showed similar, if not identical cascade dynamics. Nor is this effect reducible to postural stability, as the growth of lognormality in postural fluctuations with timescale did not depend on the viewing condition. Instead, these effects reflect endogenous postural control, as known indices of endogenous postural fluctuations moderated the effect of viewing condition on postural cascade dynamics. In short, the moderating of postural cascade dynamics by visual effort is an endogenous phenomenon that is difficult to counteract.

Past work has shown that variability in postural fluctuations is reduced when visually fixating at near as opposed to distant targets [12,13,22], an effect that has been directly attributed to the precision demands of viewing distance [32]. The present finding significantly adds to this work by showing that this effect is not linear. Visually fixating at too-close and too-far targets may introduce ambiguity between sway and optic flow: too-close targets making head-sway more likely to destabilize the posture, and too-far targets might recruit more torso-sway. However, visually fixating at an optimally placed target may resolve this ambiguity as the body courses with fluctuations ferrying information across the body. Furthermore, the stark similarities in the cascade dynamics between standing still and visually fixating at an optimal distance and standing still with eyes closed adds a novel dimension to the family of perception-action couplings in postural control. Postural control does not depend on vision per se. Instead, it depends on how visual information is laced together with activities at multiple scales: subtle fluctuations of the head, torso, and the whole body and photoreceptor support of the target’s retinal image.

Postural cascade dynamics is rooted in the known physiology. Indeed, the biophysical substrate supporting postural control is imagined as a neurally-tunable bodywide multifractal tensegrity (MFT) network consisting of the connective-tissue net and extra-cellular matrix (ECM) in which components hang together by tensional and compression forces at multiple scales, from micro (e.g., cells and membranes) to macro (e.g., muscles and bones) [43–45]. This network cooperates directly and uninterruptedly with the nervous system and exhibits similar cascade-like dynamics as in “neuronal avalanches” [46,47]. It is no small coincidence that tensegrity and avalanche models all converge on multiplicativity [48,49] that inspired multiscale quantification of non-Gaussianity [27–29]. Hence, MFT approaches suggest multiple points of entry through which stimulation elicits cascades to spread across scales— from the surface underfoot [8] to the loads at hand [9] and now from the incoming visual information. As the cascade metaphor entails, multiple scales are not merely coexisting but explicitly interacting, with influences of one spreading to others. If mechanically and visually perturbing posture engender cascade dynamics similarly [8,9], then postural control may emerge from the perceptuomotor sensitivity of specifically multiplicative fluctuations to the environment. The methods introduced here would enable future investigations into how different suprapostural tasks lace together deep texture at very many scales.

In short, the present findings suggest that visual perception by no means is a passive function proceeding in parallel to still posture—the poise carried by the body as it stands and looks suggests that moving visual targets nearer or farther brings a reorganization of postural interactions across scales. This standing-and-looking preamble to action brims with dynamic redistributions of tensional and compression forces. Indeed, the participants completed no task beyond standing still and maintaining visual fixation. But these cascades affecting a dynamic balance of forces across the body seem to be essential foundations for how the body reaches out beyond itself to engage with the environment [7,50–52]. So, the present findings show how vision plays not only upon the photoreceptors in the eyes but also across the full-body postural system.

## Supplementary material

**Table S1**. Coefficients of linear mixed-effect (LME) model examining the effect of Viewing condition on 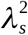 vs. log-timescale curves.

**Table S2**. Coefficients of LME model examining the effects of Viewing condition on 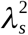 vs. log-timescale curves for orthogonal linear and quadratic polynomials, for interactions with indices of endogenous postural fluctuations.

**Table S3**. Coefficients of LME model examining the effects of Viewing condition on 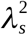 vs. log-timescale curves for orthogonal linear, quadratic, and cubic polynomials, for interactions with indices of endogenous postural fluctuations.

**Table S4**. Coefficients of generalized linear mixed-effect (GLME) model examining the effect of Viewing condition on the growth of lognormality in postural fluctuations with log-timescale.

---

**Fig. S1.**
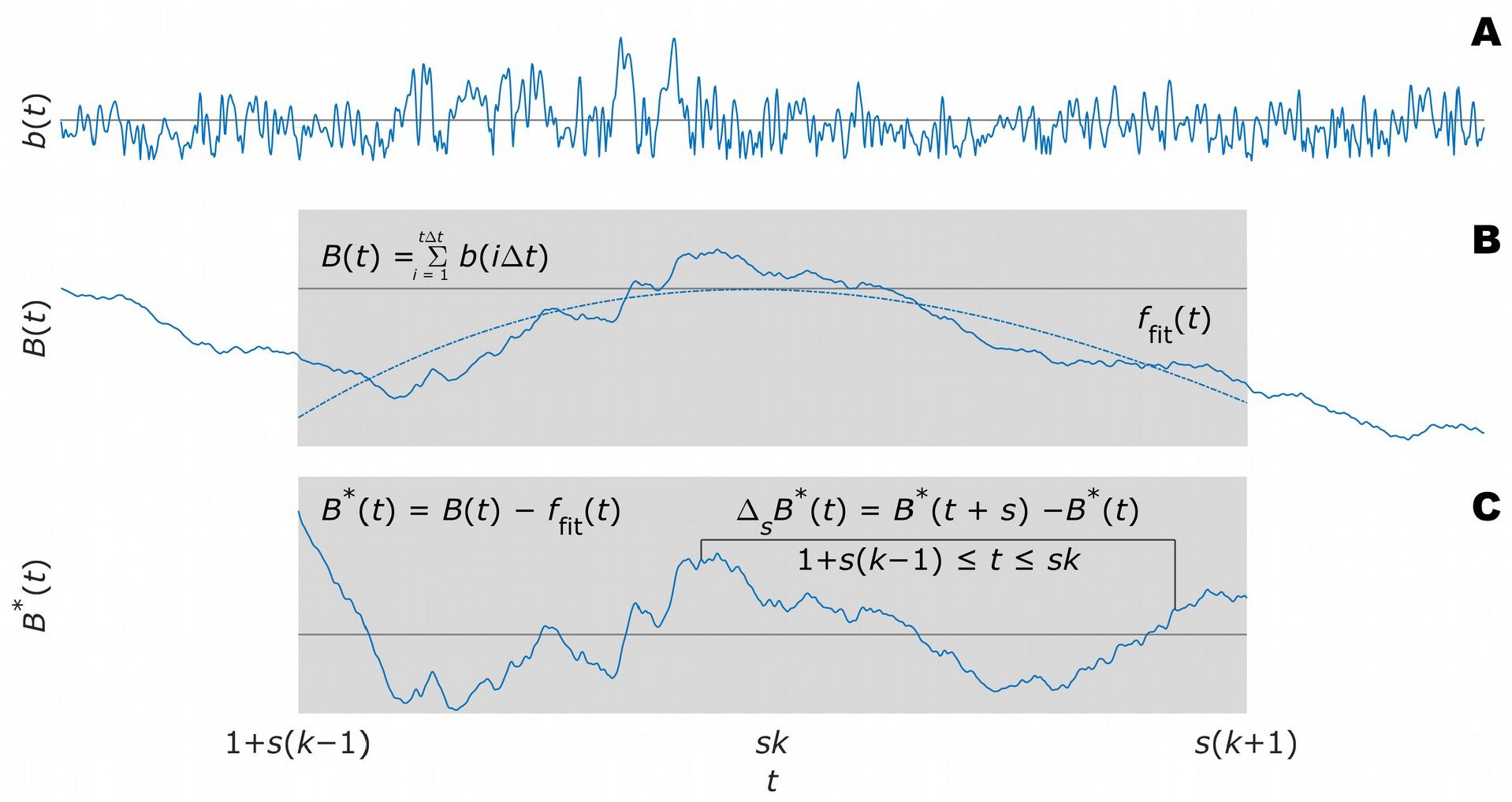
Schematic illustration of multiscale probability density function (PDF) analysis. (A) The first step is to generate {*B* (*t*)} by integrating {*b*(*t*)} after centering by the mean 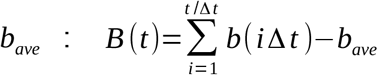. (B) A a 3^rd^ order polynomial detrends {*B* (*t*)} within *k* overlapping windows of length 2 *s, s* being the timescale. (C) Intermittent deviation Δ_*s*_ *B*(*t*) in *k*^*th*^ window from 1+*s* (*k* −1) to *sk* in the detrended time series {*B*^*d*^(*t*)=*B* (*t*)−*f* _*fit*_(*t*)} is computed as Δ_*s*_ *B*^*d*^ (*t*)=*B*^*d*^ (*t* + *s*)−*B*^*d*^(*t*), where 1+*s* (*k* −1)≤*t* ≤*sk* and *f* _*fit*_ (*t*) is the polynomial representing the local trend of {*B* (*t*)}, of which the elimination assures the zero-mean probability density function in the next step.

**Fig. S2.**
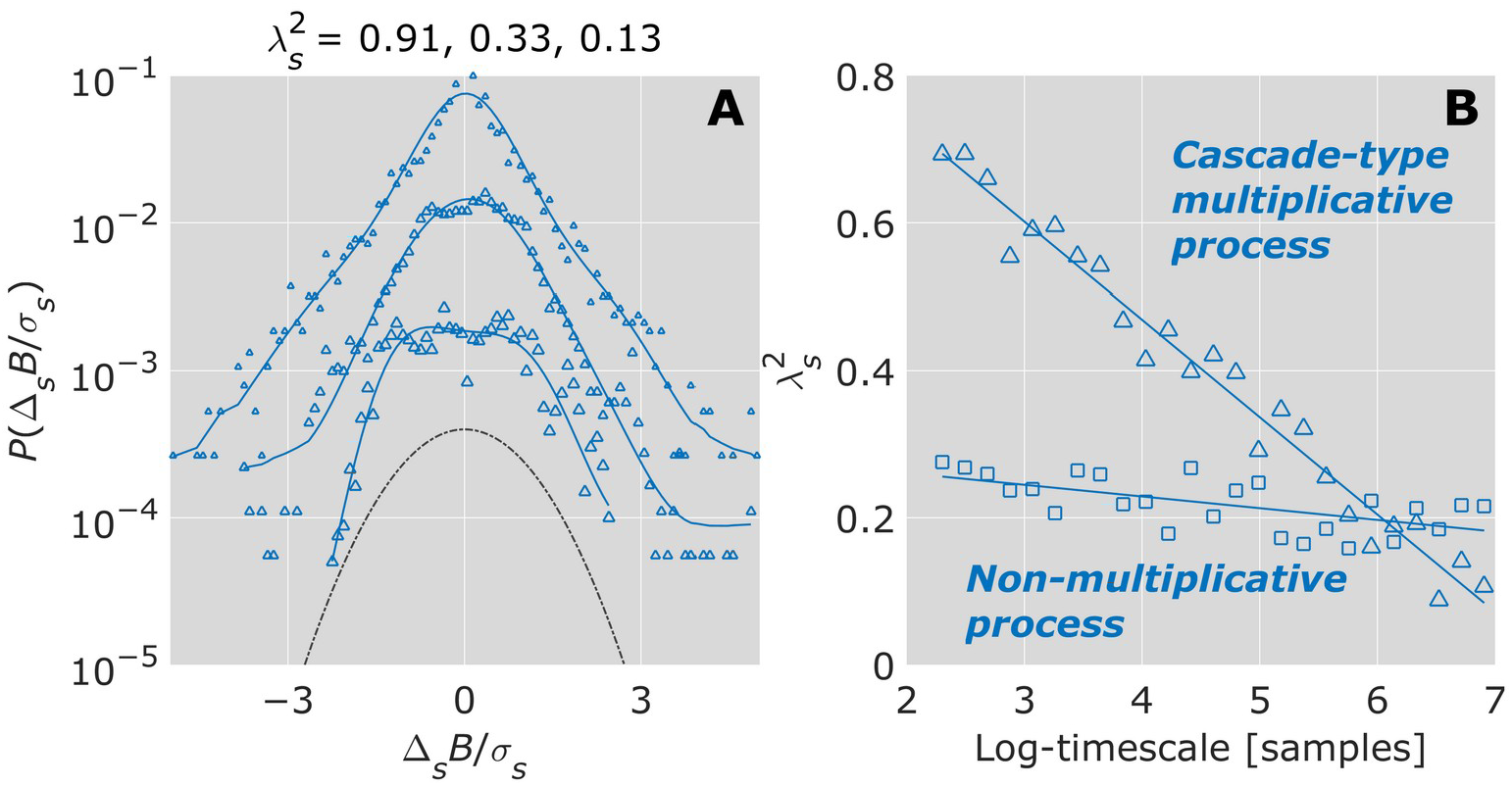
Schematic illustration of non-Gaussianity index *λ*_*s*_. (A) The relationship between 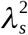 and shapes of PDF plotted in linear-log coordinates for 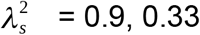, and 0.13 (from top to bottom). For clarity, the PDFs have been shifted vertically. As 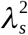 increases, the PDF becomes increasingly peaked and fat-tailed. As *λ*_*s*_ decreases, the PDF increasingly resembles the Gaussian (dashed line), assuming a perfect Gaussian as 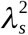 approaches 0. (B) Cascade-type multiplicative processes yield the inverse relationship 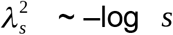.

## References

[1] J.S. Matthis, K.S. Muller, K. Bonnen, M.M. Hayhoe, Retinal optic flow during natural locomotion, BioRxiv. (2020) 2020.07.23.217893. https://doi.org/10.1101/2020.07.23.217893.

[2] J.K. O’Regan, Solving the “real” mysteries of visual perception: The world as an outside memory., Can. J. Psychol. Can. Psychol. 46 (46) 461–488. https://doi.org/10.1037/h0084327.

[3] A.S. Mauss, A. Borst, Optic flow-based course control in insects, Curr. Opin. Neurobiol. 60 (60) 21–27. https://doi.org/10.1016/j.conb.2019.10.007.

[4] B. Manor, M.D. Costa, K. Hu, E. Newton, O. Starobinets, H.G. Kang, C.K. Peng, V. Novak, L.A. Lipsitz, Physiological complexity and system adaptability: Evidence from postural control dynamics of older adults, J. Appl. Physiol. 109 (109) 1786–1791. https://doi.org/10.1152/japplphysiol.00390.2010.

[5] D.G. Kelty-Stephen, K. Palatinus, E. Saltzman, J.A. Dixon, A tutorial on multifractality, cascades, and interactivity for empirical time series in ecological science, Ecol. Psychol. 25 (25) 1–62. https://doi.org/10.1080/10407413.2013.753804.

[6] M. Mangalam, D.G. Kelty-Stephen, Multiplicative-cascade dynamics supports whole-body coordination for perception via effortful touch, Hum. Mov. Sci. 70 (70) 102595. https://doi.org/10.1016/j.humov.2020.102595.

[7] M. Mangalam, N.S. Carver, D.G. Kelty-Stephen, Multifractal signatures of perceptual processing on anatomical sleeves of the human body, J. R. Soc. Interface. 17 (17) 20200328. https://doi.org/10.1098/rsif.2020.0328.

[8] M. Mangalam, D.G. Kelty-Stephen, Hypothetical control of postural sway, BioRxiv. (2020) 104760. https://doi.org/10.1101/2020.05.19.104760.

[9] P.F. Mariusz, M. Mangalam, D.G. Kelty-Stephen, G. Juras, Postural instability recruits shorter-timescale processes into the non-Gaussian cascade processes, BioRxiv. (2020) 136895. https://doi.org/10.1101/2020.06.05.136895.

[10] M. Costa, A.L. Goldberger, C.-K. Peng, Multiscale entropy analysis of complex physiologic time series, Phys. Rev. Lett. 89 (89) 68102. https://doi.org/10.1103/PhysRevLett.89.068102.

[11] M. Costa, A.L. Goldberger, C.-K. Peng, Multiscale entropy analysis of biological signals, Phys. Rev. E. 71 (71) 21906. https://doi.org/10.1103/PhysRevE.71.021906.

[12] T.A. Stoffregen, R.J. Pagulayan, B.G. Bardy, L.J. Hettinger, Modulating postural control to facilitate visual performance, Hum. Mov. Sci. 19 (19) 203–220. https://doi.org/10.1016/S0167-9457(00)00009-9.

[13] T.A. Stoffregen, L.J. Smart, B.G. Bardy, R.J. Pagulayan, Postural stabilization of looking, J. Exp. Psychol. Hum. Percept. Perform. 25 (25) 1641–1658. https://doi.org/10.1037/0096-1523.25.6.1641.

[14] M.C. Dault, A.C.H. Geurts, T.W. Mulder, J. Duysens, Postural control and cognitive task performance in healthy participants while balancing on different support-surface configurations, Gait Posture. 14 (14) 248–255. https://doi.org/10.1016/S0966-6362(01)00130-8.

[15] O. Oullier, B.G. Bardy, T.A. Stoffregen, R.J. Bootsma, Postural coordination in looking and tracking tasks, Hum. Mov. Sci. 21 (21) 147–167. https://doi.org/10.1016/S0167-9457(02)00093-3.

[16] M.R. Giveans, K. Yoshida, B. Bardy, M. Riley, T.A. Stoffregen, Postural sway and the amplitude of horizontal eye movements, Ecol. Psychol. 23 (23) 247–266. https://doi.org/10.1080/10407413.2011.617215.

[17] T.A. Stoffregen, B.G. Bardy, C.T. Bonnet, R.J. Pagulayan, Postural stabilization of visually guided eye movements, Ecol. Psychol. 18 (18) 191–222. https://doi.org/10.1207/s15326969eco1803_3.

[18] C.M. Fiorelli, P.F. Polastri, S.T. Rodrigues, A.M. Baptista, T. Penedo, V.A.I. Pereira, L. Simieli, F.A. Barbieri, Gaze position interferes in body sway in young adults, Neurosci. Lett. 660 (660) 130–134. https://doi.org/10.1016/j.neulet.2017.09.008.

[19] T.-T. Lê, Z. Kapoula, Role of ocular convergence in the Romberg quotient, Gait Posture. 27 (27) 493–500. https://doi.org/10.1016/j.gaitpost.2007.06.003.

[20] M.C. Musolino, P.J. Loughlin, P.J. Sparto, M.S. Redfern, Spectrally similar periodic and non-periodic optic flows evoke different postural sway responses, Gait Posture. 23 (23) 180–188. https://doi.org/10.1016/j.gaitpost.2005.02.008.

[21] T.A. Stoffregen, P. Hove, J. Schmit, B.G. Bardy, Voluntary and involuntary postural responses to imposed optic flow, Motor Control. 10 (10) 24–33. https://doi.org/10.1123/mcj.10.1.24.

[22] Z. Kapoula, T.-T. Lê, Effects of distance and gaze position on postural stability in young and old subjects, Exp. Brain Res. 173 (173) 438–445. https://doi.org/10.1007/s00221-006-0382-1.

[23] J. Munafo, C. Curry, M.G. Wade, T.A. Stoffregen, The distance of visual targets affects the spatial magnitude and multifractal scaling of standing body sway in younger and older adults, Exp. Brain Res. 234 (234) 2721–2730. https://doi.org/10.1007/s00221-016-4676-7.

[24] E.A. Codling, M.J. Plank, S. Benhamou, Random walk models in biology, J. R. Soc. Interface. 5 (5) 813–834. https://doi.org/10.1098/rsif.2008.0014.

[25] M.A. Lomholt, K. Tal, R. Metzler, K. Joseph, Lévy strategies in intermittent search processes are advantageous, Proc. Natl. Acad. Sci. 105 (105) 11055–11059. https://doi.org/10.1073/pnas.0803117105.

[26] B. Castaing, Y. Gagne, E.J. Hopfinger, Velocity probability density functions of high Reynolds number turbulence, Phys. D Nonlinear Phenom. 46 (46) 177–200. https://doi.org/10.1016/0167-2789(90)90035-N.

[27] B.J. West, M. Turalska, Hypothetical control of heart rate variability, Front. Physiol. 10 (10) 1078. https://doi.org/10.3389/fphys.2019.01078.

[28] K. Kiyono, Z.R. Struzik, N. Aoyagi, S. Sakata, J. Hayano, Y. Yamamoto, Critical scale invariance in a healthy human heart rate, Phys. Rev. Lett. 93 (93) 178103. https://doi.org/10.1103/PhysRevLett.93.178103.

[29] K. Kiyono, J. Hayano, E. Watanabe, Z.R. Struzik, Y. Yamamoto, Non-Gaussian heart rate as an independent predictor of mortality in patients with chronic heart failure, Hear. Rhythm. 5 (5) 261–268. https://doi.org/10.1016/j.hrthm.2007.10.030.

[30] K. Kiyono, J. Hayano, S. Kwak, E. Watanabe, Y. Yamamoto, Non-gaussianity of low frequency heart rate variability and sympathetic activation: Lack of increases in multiple system atrophy and parkinson disease, Front. Physiol. 3 (3) 34. https://doi.org/10.3389/fphys.2012.00034.

[31] J. Hayano, K. Kiyono, Z. Struzik, Y. Yamamoto, E. Watanabe, P. Stein, L. Watkins, J. Blumenthal, R. Carney, Increased non-Gaussianity of heart rate variability predicts cardiac mortality after an acute myocardial infarction, Front. Physiol. 2 (2) 65. https://doi.org/10.3389/fphys.2011.00065.

[32] I.-C. Lee, M.M. Pacheco, K.M. Newell, The precision demands of viewing distance modulate postural coordination and control, Hum. Mov. Sci. 66 (66) 425–439. https://doi.org/10.1016/j.humov.2019.05.019.

[33] T. Schreiber, A. Schmitz, Improved surrogate data for nonlinearity tests, Phys. Rev. Lett. 77 (77) 635–638. https://doi.org/10.1103/PhysRevLett.77.635.

[34] J.S. Richman, J.R. Moorman, Physiological time-series analysis using approximate entropy and sample entropy, Am. J. Physiol. Circ. Physiol. 278 (278) H2039–H2049. https://doi.org/10.1152/ajpheart.2000.278.6.H2039.

[35] J.-H. Ko, K.M. Newell, Aging and the complexity of center of pressure in static and dynamic postural tasks, Neurosci. Lett. 610 (610) 104–109. https://doi.org/10.1016/j.neulet.2015.10.069.

[36] C.-K. Peng, S. V Buldyrev, S. Havlin, M. Simons, H.E. Stanley, A.L. Goldberger, Mosaic organization of DNA nucleotides, Phys. Rev. E. 49 (49) 1685–1689. https://doi.org/10.1103/PhysRevE.49.1685.

[37] M. Mangalam, R. Chen, T.R. McHugh, T. Singh, D.G. Kelty-Stephen, Bodywide fluctuations support manual exploration: Fractal fluctuations in posture predict perception of heaviness and length via effortful touch by the hand, Hum. Mov. Sci. 69 (69) 102543. https://doi.org/10.1016/j.humov.2019.102543.

[38] A. Chhabra, R. V Jensen, Direct determination of the f(α) singularity spectrum, Phys. Rev. Lett. 62 (62) 1327–1330. https://doi.org/10.1103/PhysRevLett.62.1327.

[39] K. Kiyono, Z.R. Struzik, Y. Yamamoto, Estimator of a non-Gaussian parameter in multiplicative log-normal models, Phys. Rev. E. 76 (76) 41113. https://doi.org/10.1103/PhysRevE.76.041113.

[40] D. Bates, M. Mächler, B. Bolker, S. Walker, Fitting linear mixed-effects models using lme4, ArXiv. (2014) 5823. https://doi.org/arXiv:1406.5823.

[41] A. Kuznetsova, P.B. Brockhoff, R.H.B. Christensen, lmerTest package: Tests in linear mixed effects models, J. Stat. Softw. 82 (82) 1548–7660. https://doi.org//10.18637/jss.v082.i13.

[42] W. Lee, K.J. Grimm, Generalized linear mixed-effects modeling programs in R for binary outcomes, Struct. Equ. Model. A Multidiscip. J. 25 (25) 824–828. https://doi.org/10.1080/10705511.2018.1500141.

[43] R. Schleip, F. Mechsner, A. Zorn, W. Klingler, The bodywide fascial network as a sensory organ for haptic perception, J. Mot. Behav. 46 (46) 191–193. https://doi.org/10.1080/00222895.2014.880306.

[44] M.T. Turvey, S.T. Fonseca, The medium of haptic perception: A tensegrity hypothesis, J. Mot. Behav. 46 (46) 143–187. https://doi.org/10.1080/00222895.2013.798252.

[45] D.E. Ingber, From cellular mechanotransduction to biologically inspired engineering, Ann. Biomed. Eng. 38 (38) 1148–1161. https://doi.org/10.1007/s10439-010-9946-0.

[46] S.R. Miller, S. Yu, D. Plenz, The scale-invariant, temporal profile of neuronal avalanches in relation to cortical γ–oscillations, Sci. Rep. 9 (9) 16403. https://doi.org/10.1038/s41598-019-52326-y.

[47] D.R. Chialvo, Emergent complex neural dynamics, Nat. Phys. 6 (6) 744–750. https://doi.org/10.1038/nphys1803.

[48] T. Zorick, J. Landers, A. Leuchter, M.A. Mandelkern, EEG multifractal analysis correlates with cognitive testing scores and clinical staging in mild cognitive impairment, J. Clin. Neurosci. 76 (76) 195–200. https://doi.org/10.1016/j.jocn.2020.04.003.

[49] A. Das, A. Levina, Critical neuronal models with relaxed timescale separation, Phys. Rev. X. 9 (9) 21062. https://doi.org/10.1103/PhysRevX.9.021062.

[50] M. Mangalam, N.S. Carver, D.G. Kelty-Stephen, Global broadcasting of local fractal fluctuations in a bodywide distributed system supports perception via effortful touch, Chaos, Solitons & Fractals. 135 (135) 109740. https://doi.org/10.1016/j.chaos.2020.109740.

[51] N.S. Carver, D. Bojovic, D.G. Kelty-Stephen, Multifractal foundations of visually-guided aiming and adaptation to prismatic perturbation, Hum. Mov. Sci. 55 (55) 61–72. https://doi.org/10.1016/j.humov.2017.07.005.

[52] C. Bell, N. Carver, J. Zbaracki, D. Kelty-Stephen, Nonlinear amplification of variability through interaction across scales supports greater accuracy in manual aiming: Evidence from a multifractal analysis with comparisons to linear surrogates in the Fitts task, Front. Physiol. 10 (10) 998. https://doi.org/10.3389/fphys.2019.00998.

